# Rac1-Abi-1 Signaling Links DNA Repair to Tumorigenesis

**DOI:** 10.64898/2026.11.13.766481

**Authors:** Yiheng Huang, Jie Xu, Ziqing Zhu, Angelica Lin, Sumeet Solanki, Kari Wilder-Romans, Jihan Park, Jing Li, Honglai Zhang, Timothy Jho, Daven Kothari, Wilson Zheng, Yuezhong Zhang, Elizabeth McCulla, Mingjia Tan, Theodore S. Lawrence, Meredith A. Morgan, Menggang Yu, Benjamin E Curtis, Thomas L. Saunders, Zachary T. Freeman, Sriram Venneti, Yatrik M. Shah, Daniel R. Wahl, Weihua Zhou

## Abstract

The DNA damage response governs genomic integrity, and abnormal DNA repair can drive tumorigenesis. We previously identified a Rac1-Abi-1 signaling pathway in which Rac1 activates protein phosphatase 5, leading to Abi-1-S323 dephosphorylation and enhanced non-homologous end joining (NHEJ). Given the error-prone nature of NHEJ and its potential to promote genomic instability, we sought to determine whether Rac1-Abi-1-mediated regulation of NHEJ contributes to tumorigenesis. Here, we demonstrate the tumorigenic role of Abi-1-S323 dephosphorylation in nonmalignant cell models and multiple tumorigenesis models established using a newly generated genetically engineered Abi-1 mouse model. We first found that the dephospho-mimetic Abi-1-S323A mutant promoted transformation-associated phenotypes in nonmalignant human cells and enhanced orthotopic glioblastoma growth under unperturbed conditions. To further investigate its tumorigenic potential *in vivo*, we generated CRISPR/Cas9-mediated Abi-1-S296A knock-in mice, in which S296 corresponds to human S323. Abi-1-S296A mouse embryonic fibroblasts exhibited increased proliferation, while mutant mice developed spontaneous tumors involving lymphoid tissues and organs under unperturbed conditions. Furthermore, Abi-1-S296A increased susceptibility to tumorigenesis following DNA damage. In a low-dose whole-body irradiation model, Abi-1-S296A mice showed a trend toward shortened survival and increased tumor development, with lesions including Harderian gland papillary cystadenomas, round-cell neoplasms, and anaplastic sarcomas. In addition, in an azoxymethane/dextran sulfate sodium (AOM/DSS)-induced colorectal tumor model, Abi-1-S296A mice developed significantly more tumors and had a greater tumor burden. These findings identify Abi-1-S323 dephosphorylation as a previously unrecognized regulator of tumorigenesis and establish a new mouse model for investigating the physiological and pathological consequences of constitutive activation of upstream NHEJ-regulatory signaling.

## Introduction

The DNA damage response (DDR) serves as a critical safeguard against genomic instability. DNA damage and its aberrant repair contribute to the development of many cancers [1]. Among various forms of DNA damage, DNA double-strand breaks (DSBs) are particularly cytotoxic [2]. They can arise from exogenous sources, such as ionizing radiation and chemical agents, as well as endogenous processes, including oxidative stress, replication stress, and metabolic disturbances [3]. To maintain genomic integrity, cells have evolved multiple pathways to repair DSBs, primarily homologous recombination (HR) and non-homologous end joining (NHEJ)[4]. HR repairs DSBs by using a homologous DNA sequence as a template to guide accurate repair, whereas NHEJ directly rejoins broken DNA ends without requiring a homologous template and can be error prone. Improper DSB repair can lead to chromosomal aberrations, thereby promoting genomic instability and tumorigenesis [5, 6].

Although essential for DSB repair, the error-prone nature of NHEJ can promote genomic instability and tumorigenesis [7]. Aberrant activation or dysregulation of NHEJ components has been reported in numerous malignancies, including prostate [8], colon [9, 10], lung [11], liver [12, 13], esophagus [14], and gastric cancers [15]. Furthermore, pharmacological inhibition of NHEJ has been shown to suppress tumor progression in preclinical models [16]. Although previous studies have investigated how NHEJ factors or components contribute to tumorigenesis, it remains unclear whether upstream DDR signaling pathways promote tumorigenesis by regulating NHEJ.

We previously identified a DDR signaling pathway in which increased GTP levels activate Rac1, leading to activation of protein phosphatase 5 (PP5) and subsequent dephosphorylation of serine 323 on Abl-interactor 1 (Abi-1), thereby promoting NHEJ and radiation resistance in glioblastoma [17]. More recently, we found that this signaling pathway regulates NHEJ and radiation resistance across multiple cancer types [18]. However, whether Rac1-Abi-1 signaling also contributes to tumorigenesis through its regulation of NHEJ remains unknown.

Here, we investigated the tumorigenic potential of Rac1-Abi-1 signaling using nonmalignant cellular models, an orthotopic GBM model, and multiple tumorigenesis models established using a newly generated CRISPR/Cas9-mediated Abi-1-S296A knock-in mouse model, which carries an alanine substitution at the murine residue corresponding to human Abi-1-S323 and constitutively maintains the signaling activation. We found that expression of the dephospho-mimetic Abi-1-S323A mutant promoted transformation-associated phenotypes in nonmalignant human cells and enhanced glioblastoma growth under unperturbed conditions. In the Abi-1-S296A knock-in mouse model, we observed spontaneous tumor development under unperturbed conditions, broadly increased tumor development across multiple tissues following DNA damage induced by low-dose whole-body irradiation, and enhanced colon tumorigenesis in an azoxymethane/dextran sulfate sodium (AOM/DSS) model that combines AOM-induced DNA damage with DSS-induced colitis. These findings establish Abi-1-S323 dephosphorylation as a previously unrecognized driver of tumorigenesis and reveal an important role for Rac1-Abi-1 signaling in cancer development.

### Methods and Material

### Cells and Reagents

Cells were maintained at 37°C in a humidified incubator with 5% CO₂ in medium supplemented with 10% fetal bovine serum (FBS; Cat# A5256801, Gibco), 100 μg/mL Normocin (Cat# ant-nr-1, InvivoGen), and 100 U/mL penicillin–streptomycin–glutamine (Cat# 10378-016, Gibco). The immortalized normal human bronchial epithelial cell line BEAS-2B was obtained from the American Type Culture Collection (ATCC) and cultured in Airway Cell Basal Medium (PCS-300-030, ATCC) [19], whereas NCM460 cells were cultured in Dulbecco’s modified Eagle medium (DMEM; Cat# 11965-092, Gibco) [20]. The GBM38 cell line, previously described in our studies [17, 21], and normal human astrocytes, provided by Dr. Sofia Merajver (University of Michigan) [18, 22], were cultured in DMEM. Mycoplasma testing was performed monthly using the MycoAlert™ Mycoplasma Detection Kit (Cat# LT07-418, Lonza).

### Generation and culture of mouse embryonic fibroblasts

Primary mouse embryonic fibroblasts (MEFs) were isolated from individual embryos at ∼E13.5 as described previously [23]. Briefly, embryos were collected and rinsed with PBS followed by DMEM supplemented with 25 mM HEPES. Embryonic tissues were minced into small fragments and enzymatically digested with 0.05% trypsin containing 0.53 mM EDTA for 20 min at 37°C. Residual tissue clumps were further dissociated by passing the suspension three times through an 18-gauge needle. The resulting cell suspensions were collected and plated in MEF culture medium consisting of 15% FBS, 2 mM L-glutamine, 0.1 mM MEM non-essential amino acids, and 10 μg/mL gentamicin. Cells were maintained at 37°C in a humidified incubator with 5% CO₂.

### Immunoblotting assay

Human cells or MEFs were lysed in RIPA lysis buffer (Cat# 89900, Thermo Scientific) supplemented with PhosSTOP phosphatase inhibitor (Cat# 04906845001, Roche) and complete protease inhibitor tablets (Cat# 1187358001, Roche). Proteins were detected using primary antibodies at a 1:1,000 dilution, including an anti-β-actin antibody (Cat# sc47778, Santa Cruz Biotechnology) and a rabbit polyclonal anti-p-Abi-1 (Ser323) antibody generated by Cell Signaling Technology, as described previously [17].

### Soft agar colony formation assay

A standard soft agar assay was performed as described previously [24]. Briefly, BEAS-2B cells transfected with Abi-1 wild-type, S323A, or S323D constructs were suspended in 0.33% agar containing 10% FCS and plated in 60-mm dishes. Cells were cultured at 37°C for 14 days and then stained overnight with p-iodonitrotetrazolium (1 mg/mL; Sigma). Colonies containing more than eight cells were counted in five randomly selected fields per dish. Results were obtained from three independent experiments.

### Celltiter-Glo cell viability assay

Normal human astrocytes or NCM460 normal human colon epithelial cells were transfected with the indicated Abi-1 plasmids and seeded into 96-well plates at approximately 2,000 cells per well. Cell viability was measured daily over a total period of 5 or 7 days using the CellTiter-Glo® 2D Cell Viability Assay (Cat# G9242, Promega), according to the manufacturer’s instructions.

### Long-term neurosphere assay and live cell imaging

The long-term neurosphere assay with automated imaging was performed using the BioSpa and Cytation systems as described previously [25]. Briefly, GBM38 stable cell lines expressing Abi-1-WT, S323D, or S323A were replated at low density (500 cells per well) in 96-well plates. The plates were incubated for 8 days in a BioSpa incubator (Agilent Technologies) and imaged every 24 h using a Cytation 5 cell imaging reader equipped with a 4× objective. Images were analyzed using Gen5 software (Agilent Technologies). A two-dimensional area threshold of 2,600 μm² was applied to identify neurospheres and the number of neurospheres was quantified at the endpoint.

### Orthotopic glioma models

GBM38 orthotopic models were established as previously described [17]. All animal experiments were approved by the University Committee on Use and Care of Animals (UCUCA) at the University of Michigan. Briefly, luciferase-expressing GBM38 stable cell lines expressing the Abi-1-S323D or Abi-1-S323A mutant were intracranially injected into 8–10-week-old Rag1-knockout C57BL/6 mice (Strain #002216). Postoperative care was provided for 7–10 days. Tumor growth was monitored by bioluminescence imaging using an IVIS™ Spectrum imaging system (PerkinElmer). Mouse survival was monitored until animals reached humane endpoints, as defined by institutional guidelines.

### Generation of Abi1 S296A knock-in mice

CRISPR/Cas9 technology was used to introduce mutations upstream of Abi1 (MGI: 1913860; ENSMUSG00000018381). The CRISPOR algorithm [26] was used to identify a single guide RNA target (sgRNA) predicted to cut the chromosome. sgRNA C386C, used for the 5′ chromosome break, targets the forward strand with the protospacer sequence 5′-CCCCAAGTGACAGGACAGAG -3′ and PAM GGG. sgRNA C386X, used for the 5′ chromosome break, targets the reverse strand with the protospacer sequence 5′-TTGCAGGCATACAGCAATAG-3′ and PAM TGG. C386C and C386X had Cutting Frequency Determination scores of 72 and 89, Moreno-Mateos activity scores of 51 and 57, respectively [26, 27]. The sgRNA (60 ng/ul) was complexed with HiFi Cas9 protein (50 ng/ul) from IDT and individually tested to determine if ribonucleoprotein (RNP) complexes cause chromosome breaks in mouse zygotes. RNPs were microinjected into fertilized mouse eggs. Eggs were placed in culture until they developed into blastocysts. DNA was extracted from individual blastocysts for analysis. PCR with primers spanning the predicted cut site was used to generate amplicons for Sanger sequencing [28]. Amplicons were produced with C386 forward primer: 5’ CTTTATACACTTGTCCTTCATCCCCTCAG 3’ and C386 reverse primer 5’: TTTGTGTCTTCGTACCATTACTGAACCAT 3’; 1008 bp amplicon. Sanger sequencing from individual blastocysts were evaluated to determine if small insertions/deletions caused by non-homologous endjoining (NHEJ) repair of chromosome breaks were present [29]. sgRNA C386C and X were found to induce chromosome breaks.

RNPs were mixed with a spot dialyzed synthetic short single stranded DNA donor (5 ng/ul, IDT.com) prior to microinjection into mouse zygotes [30]. Silent coding changes in the sgRNA binding sequence were included in the oligonucleotide to block cutting by Cas9 after repair of the chromosome by homology directed repair [31]. The CRISPR reagents were microinjected into fertilized mouse eggs produced by mating superovulated C57BL/6:SJL (B6SJLF1) F1 Hybrids female mice (Jackson Laboratory stock no. 100012) with B6SJLF1 male mice as described [32]. CRISPR/Cas9 microinjection of zygotes produced potential founder mice which were genotyped using the following primers.

Primers used for genotyping and gRNA activity assay were forward primer: 5’ CTTTATACACTTGTCCTTCATCCCCTCAG 3’ and C386 reverse primer 5’: TTTGTGTCTTCGTACCATTACTGAACCAT 3’; 1008 bp amplicon. Amplicons were purified and Sanger sequenced. Sequenced electropherograms of amplicons from mice were evaluated to identify peaks consistent with the designed modification. To confirm the correct knock-in into *Abi1*, TOPO TA cloning was performed on DNA from animals carrying the introduced changes.

The sequence of the single stranded oligonucleotide DNA donors for Abi-1-S296A were: CTGTCTTCTTGTTTATATACTTTAGAAGTGGAGATAATTTGATTTTCATCTTCTTTTTTTATTTTATGTCTGGGCATTCCAAAAAAAATAAGAACCCCCTaCCCTCTGTCCTGTCACTTGGGGTGTTTGTAATGCTTTCTTGGTTGCCTTTGACTAATGCTTTTCAGCAGCCCCTGGCGCAGCTCCTGGTTCCCAGTATGGCACAATGACCAGGCAGATTTCTCGACACAACgCTACCACTTCTTCGACATCTTCTGGTGGATATAGACGAACTCCTTCTGTGGCCGCCCAATTCTCTGCTCAGCCTCATGTTAATGGAGGTCCACTTTATTCTCAAAATTCAAGTAAGCTTGTGTTTGCAGGCATACAGCAATAGTGtAGCTCCTTTTTAAAGGAAATTATTTGAAACTGTTATTCTTGACTGTGATAGACAACAATTTAAGCCATGCTATTTCATGCTTGTTTTTGCTTTGAATG.

### Hematoxylin and eosin (H&E) staining

Mouse tissues were fixed in 10% neutral-buffered formalin and embedded in paraffin. Paraffin-embedded tissues were sectioned at 5 µm, deparaffinized in xylene, and rehydrated through a graded ethanol series. Sections were stained with hematoxylin and eosin, followed by dehydration, clearing, and mounting.

### Whole body radiation

C57BL/6J wild type (The Jackson Laboratory; Strain #:000664) and CRISPR/Cas9 Abi-1-S296A whole body irradiation model was established as described before [33, 34]. Irradiation was performed using a Polarist Series III system (Kimtron, Oxford, CT) at the University of Michigan Comprehensive Cancer Center Experimental Irradiation Core (Ann Arbor, MI). Five-week-old mice were subjected to one fraction of 4 Gy of X-ray irradiation. Mouse survival was monitored and recorded. Mice were euthanized upon reaching predefined humane endpoints, and tissues and organs were harvested for further analyses.

### Azoxymethane (AOM)/dextran sulfate sodium (DSS)-induced colon cancer model

The AOM/DSS-induced colon cancer model in C57BL/6J wild-type and Abi-1-S296A mice was established as previously described [35]. Mice aged 8 weeks of both sexes were included. Mice received a single intraperitoneal injection of AOM (10 mg/kg; Sigma-Aldrich, A5486). Three days later, mice were given 1.5% DSS (MP Biomedicals, 160110) in their drinking water for 6 days, followed by regular drinking water for 2 weeks. This DSS treatment cycle was repeated three times. Body weight and general condition were monitored throughout the experiment. Mice were euthanized approximately 80 days after AOM administration. Colons were collected, flushed with PBS, opened longitudinally, and examined for gross tumors. Tumor numbers and dimensions were recorded. Colons were subsequently fixed in 10% neutral-buffered formalin and embedded in paraffin for histological evaluation.

### Statistics

Statistical analyses of soft agar assay results and mouse tumor numbers and volumes in AOM/DSS model were performed using two-tailed Student’s t-tests in GraphPad Prism version 11. CellTiter-Glo cell proliferation and Cytation-based neurosphere formation assay data were analyzed using two-way ANOVA. Mouse survival was analyzed using Kaplan–Meier survival curves, and differences between groups were assessed using the log-rank (Mantel–Cox) test.

## Results

### Abi-1 S323 dephosphorylation promotes normal cell transformation

We previously identified that Rac1-Abi-1 signaling promotes NHEJ repair [17]. Because NHEJ is an error-prone repair pathway that may contribute to gene mutations and deletions, thereby promoting tumorigenesis [7], we next investigated the tumorigenic potential of this signaling pathway. We first assessed cell transformation by overexpressing Abi-1-WT, the phospho-mimetic Abi-1-S323D mutant (which inhibits the signaling pathway), or the dephospho-mimetic Abi-1-S323A mutant (which activates the signaling pathway) in normal human bronchial epithelial cells (**Fig. 1A**), followed by anchorage-independent soft agar assays. Abi-1-S323D significantly suppressed, whereas Abi-1-S323A significantly increased, colony formation compared with Abi-1-WT (**Fig. 1B–C**).

**Figure 1.**
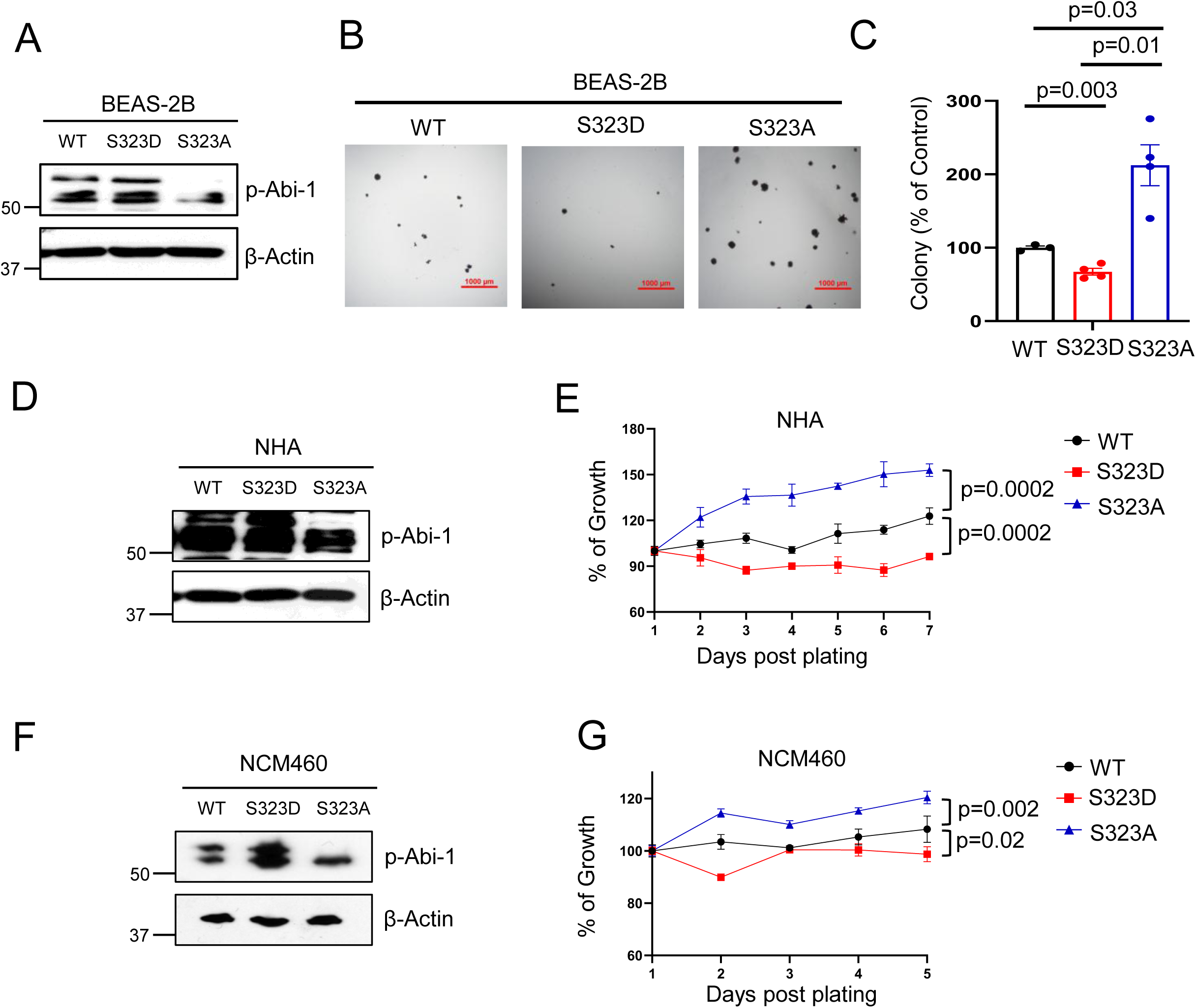
Abi-1 S323 dephosphorylation promotes normal cell transformation. **(A–C)** BEAS-2B cells were transfected with the indicated Abi-1 plasmids, followed by immunoblotting to assess protein expression (A). The remaining cells were replated in 6-cm plates for soft agar colony formation assays. After staining, colonies from 5–10 randomly selected fields were imaged (B) and counted. Quantification from three to four independent biological replicates is shown in (C). Data are presented as mean ± SEM and were analyzed using a two-tailed Student’s t-test. **(D–G)** Normal human astrocytes (NHA) and normal human colon epithelial NCM460 cells were transfected with the indicated Abi-1 plasmids, followed by immunoblotting to assess protein expression and replating in 96-well plates for CellTiter-Glo cell viability assays. Data are presented as mean ± SEM and were analyzed using two-way ANOVA.

We further investigated these effects in two additional nonmalignant human cell lines: normal human astrocytes (NHA) and normal human colonic epithelial cells (NCM460). Because neither cell line formed anchorage-independent colonies under our experimental conditions (data not shown), we performed long-term CellTiter-Glo proliferation assays to assess cell growth. Consistently, Abi-1-S323A promoted cell proliferation, whereas Abi-1-S323D inhibited proliferation compared with Abi-1-WT in both NHA (**Fig. 1D–E**) and NCM460 cells (**Fig. 1F–G**). These findings suggest that Abi-1 S323 dephosphorylation promotes cell proliferation and transformation-associated phenotypes in nonmalignant human cells.

### Abi-1 S323 dephosphorylation promotes GBM neurosphere formation and orthotopic tumor growth under unperturbed conditions

Based on the above findings demonstrating its transformation-associated phenotypes (**Fig. 1**), together with our previous finding that Abi-1 S323 dephosphorylation promotes GBM cell growth and confers radiation resistance [17], we next investigated whether this signaling pathway promotes tumor growth under unperturbed conditions. We first used a Cytation-based assay to monitor the dynamics of neurosphere formation. In patient-derived GBM38 models, forced activation of the signaling pathway by Abi-1-S323A promoted neurosphere formation, as demonstrated by increased sphere size and number, whereas inhibition of the pathway by Abi-1-S323D suppressed both phenotypes (**Fig. 2A–B**). To further evaluate the effects of Abi-1 S323 dephosphorylation in the brain microenvironment, we established orthotopic GBM models using the GBM38 stable cell line. Consistently, Abi-1-S323A promoted intracranial tumor growth (**Fig. 2C**) and shortened mouse survival (**Fig. 2D**). These findings support a role for Abi-1 S323 dephosphorylation in promoting GBM growth under unperturbed conditions and warrant further investigation in primary genetically engineered mouse models (GEMMs) to establish its role in tumorigenesis.

**Figure 2.**
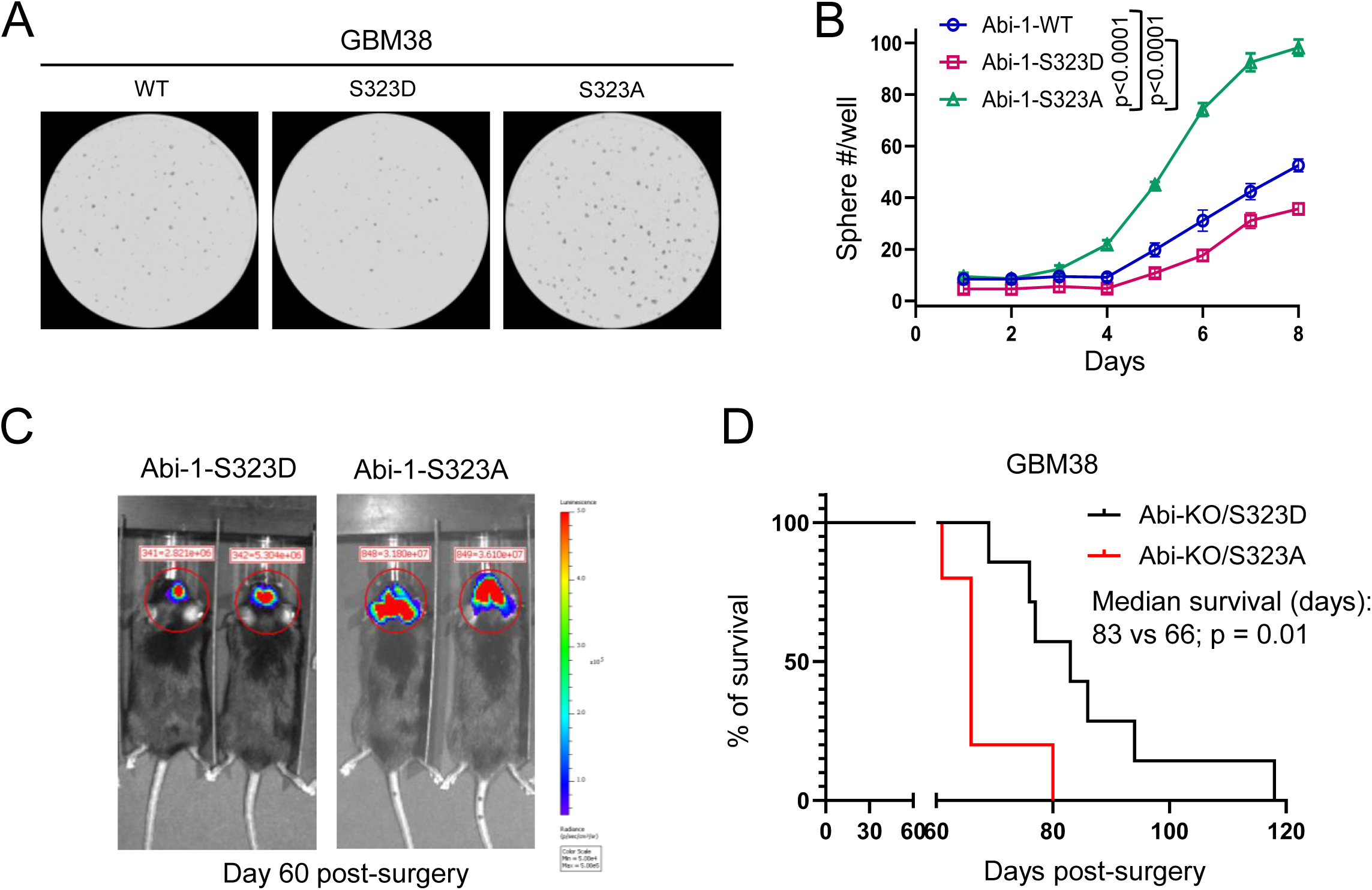
Abi-1 S323 dephosphorylation promotes GBM neurosphere formation and orthotopic tumor growth under unperturbed conditions. **(A–B)** GBM38 stable cell lines expressing Abi-1-WT, Abi-1-S323D, or Abi-1-S323A were seeded into 96-well plates and placed in a Cytation imaging system to monitor neurosphere formation. Representative images of each group are shown in (A). The number of neurospheres per well was counted, and data from six independent biological replicates were quantified in (B). Data are presented as mean ± SEM and were analyzed using two-way ANOVA. **(C)** An orthotopic GBM model was established using luciferase-expressing GBM38 cells carrying Abi-1-S323D or Abi-1-S323A. Representative bioluminescence images of mice bearing orthotopic tumors are shown. **(D)** Kaplan–Meier survival curves of mice bearing orthotopic GBM tumors. Mice were monitored daily and euthanized upon development of neurological symptoms. The p value was determined using the log-rank (Mantel–Cox) test.

### Abi-1-S296A promotes cell growth in MEFs derived from newly built CRISPR/Cas9-Abi-1-S296A knock-in mice

Conventional whole-body knockout of Abi-1 results in embryonic lethality [36], prompting the development of conditional and inducible Abi-1 knockout mouse models to investigate its functions in specific tissues and at defined developmental stages[37, 38]. However, to our knowledge, no GEMM carrying a site-specific Abi-1 mutation has been reported to date. In collaboration with the Transgenic Animal Model Core, we successfully generated a CRISPR/Cas9-mediated Abi-1-S296A knock-in mouse model, in which murine S296 corresponds to human S323 (see Methods for details). The Abi-1-S296A mice were viable, with no evidence of embryonic lethality. We validated the knock-in mutation using two independent approaches. First, Sanger sequencing of genomic DNA extracted from mouse tail tips confirmed that the murine codon 296 was changed from TCT (encoding serine, S) in WT mice to GCT (encoding alanine, A) in S296A knock-in mice (**Fig. 3A**). Second, we isolated mouse embryonic fibroblasts (MEFs) from WT and S296A embryos at approximately embryonic day 13.5 (E13.5) and performed immunoblotting using our previously developed phospho-Abi-1-S296 antibody [17]. Consistent with the sequencing results, immunoblotting confirmed the loss of Abi-1 S296 phosphorylation in S296A MEFs, as evidenced by the absence of the p-Abi-1 band compared with WT controls (**Fig. 3B**).

**Figure 3.**
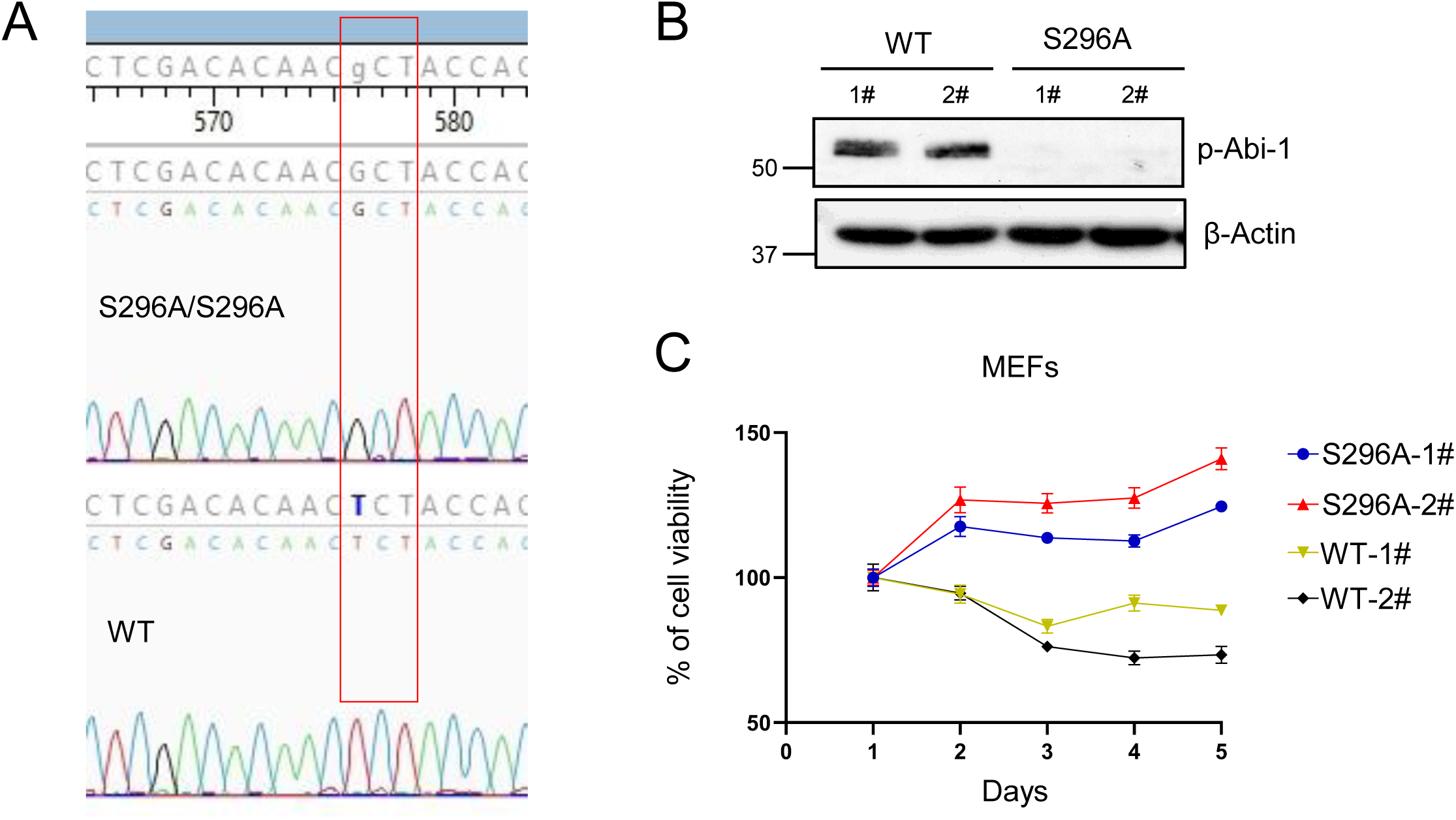
Abi-1-S296A promotes cell growth in MEFs derived from newly built CRISPR/Cas9-Abi-1-S296A knock-in mice. **(A)** Genomic DNA isolated from mouse tail biopsies was analyzed by Sanger sequencing, confirming that the murine codon 296 was changed from TCT (serine; S) in WT mice to GCT (alanine; A) in Abi-1-S296A mice. **(B)** MEFs were isolated from WT and Abi-1-S296A mice and harvested for immunoblotting using our newly generated phospho-Abi-1-S323 antibody. **(C)** Early-passage (P5) MEFs were infected with SV40 large T antigen and replated 72 hours after infection for long-term cell proliferation analysis using the CellTiter-Glo assay.

To assess cell growth, we transformed primary, non-immortalized MEFs by infecting them with oncogenic SV40 large T antigen [39] and monitored cell proliferation daily using CellTiter-Glo assays. Consistent with our findings in normal human cells, Abi-1-S296A MEFs exhibited faster growth than WT MEFs (**Fig. 3C**). These findings further support a role for Abi-1 S296 dephosphorylation in promoting cell proliferation and suggest its potential contribution to tumorigenesis.

### Abi-1-S296A promotes spontaneous tumorigenesis under unperturbed conditions

To evaluate the tumorigenic potential of Abi-1-S296A, we monitored the general health and survival of S296A mice and their WT littermate controls over an extended period (approximately 600 days). Two S296A mice were euthanized on days 180 and 420 due to moribund conditions, (**Table 1**). The remaining S296A mice were euthanized on day 579, together with the WT control mice, which remained clinically healthy throughout the observation period. Multiple organs, including the thymus, liver, spleen, lymph nodes, brain, kidneys, intestinal tract, and lungs, were collected for histopathological evaluation by H&E staining. No tumors or apparent hyperplastic lesions were identified in the examined organs of WT mice, including lymphoid tissues (**Fig. 4A**). In contrast, one S296A mouse exhibited leukemia, identified in liver and multiple tissues (**Fig. 4B**), while another showed disruption of splenic, thymic, and lymph node architecture by sheets of neoplastic round cells that expanded and effaced the normal lymphoid follicles (**Fig. 4C**). No tumors were detected in the other non-lymphoid tissues examined from the S296A mice. The characteristics of the two mouse cohorts and the observed tumor development are summarized in **Table 1**. These findings demonstrate that Abi-1 S296 dephosphorylation promotes spontaneous tumor development under unperturbed conditions.

**Figure 4.**
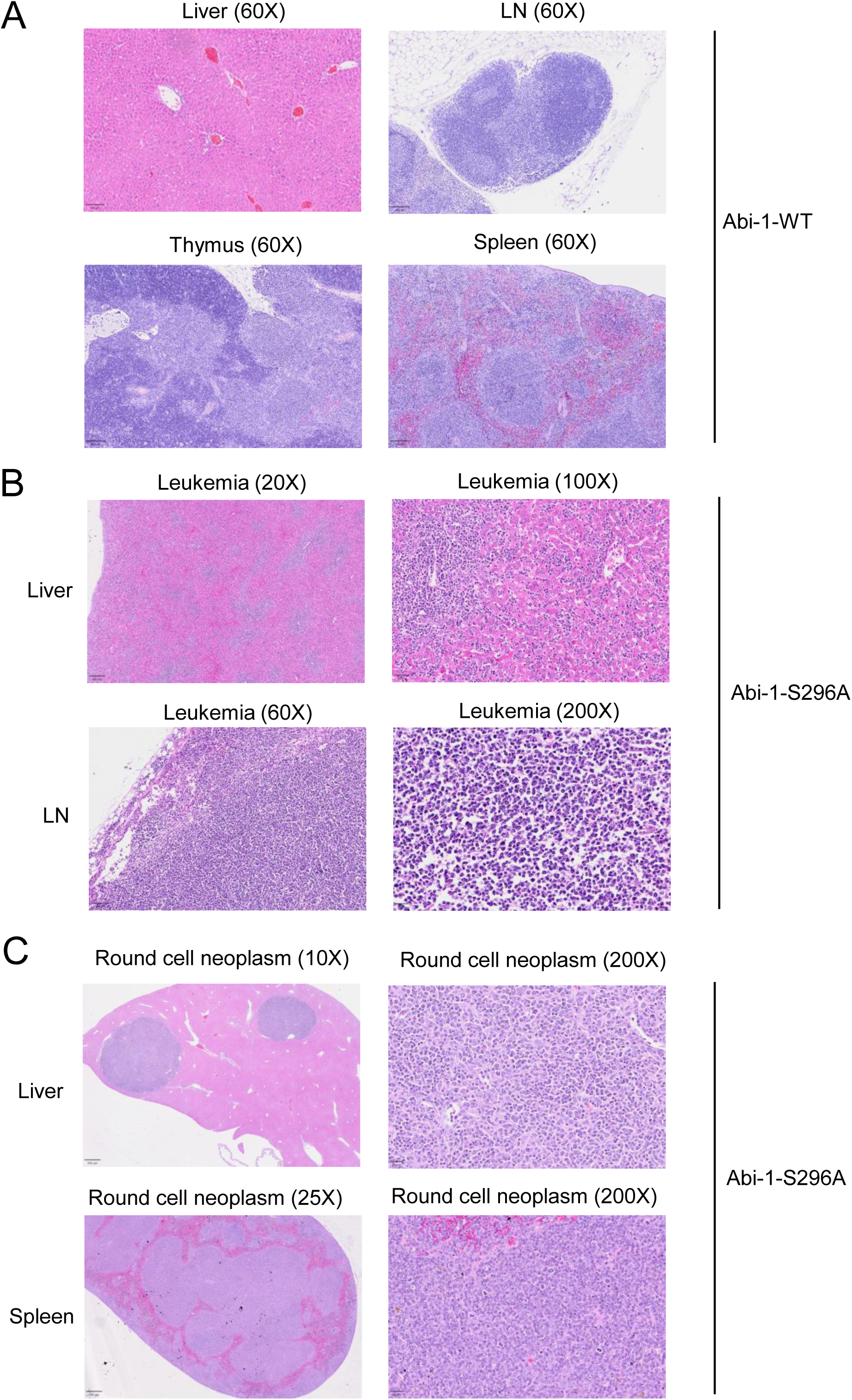
Abi-1-S296A promotes spontaneous tumorigenesis under unperturbed conditions. **(A)** Representative H&E-stained sections of the liver, lymph node (LN), thymus, and spleen from WT mice. **(B)** Representative H&E-stained sections showing leukemia involving the liver and lymph node in Abi-1-S296A mice. **(C)** Representative H&E-stained sections showing round-cell neoplasms consistent with lymphoma in the liver and spleen of Abi-1-S296A mice.

**Table 1.**
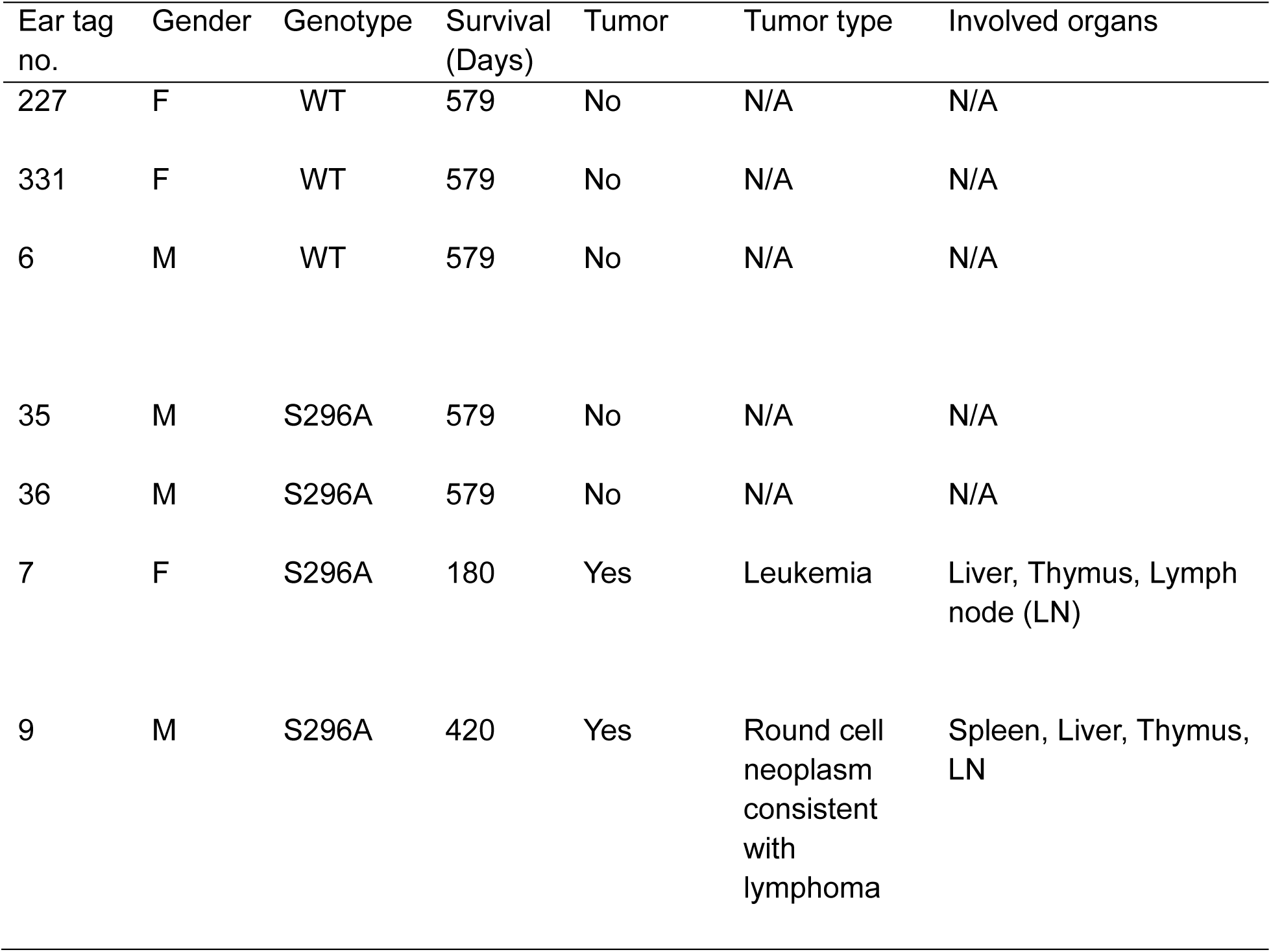
Survival Times and Tumor Type in Individual Mice under Unperturbed conditions.

| Ear tag no. | Gender | Genotype | Survival (Days) | Tumor | Tumor type | Involved organs |
|---|---|---|---|---|---|---|
| 227 | F | WT | 579 | No | N/A | N/A |
| 331 | F | WT | 579 | No | N/A | N/A |
| 6 | M | WT | 579 | No | N/A | N/A |
| 35 | M | S296A | 579 | No | N/A | N/A |
| 36 | M | S296A | 579 | No | N/A | N/A |
| 7 | F | S296A | 180 | Yes | Leukemia | Liver, Thymus, Lymph node (LN) |
| 9 | M | S296A | 420 | Yes | Round cell neoplasm consistent with lymphoma | Spleen, Liver, Thymus, LN |

### Abi-1-S296A mice show shorter survival and increased tumor development following low-dose whole-body irradiation

Since Abi-1 S323 dephosphorylation promotes NHEJ-mediated DNA repair following DNA damage, we next investigated whether the tumorigenic potential associated with this signaling state increases susceptibility to tumor development following DNA damage. Ionizing radiation is commonly used to induce DNA damage and promote tumorigenesis in mouse models [33, 34]. We therefore examined tumor development in Abi-1-S296A mice following low-dose whole-body irradiation (WBI). Five-week-old Abi-1-WT and S296A mice received a single 4-Gy dose of WBI, and their general health and survival were subsequently monitored (**Fig. 5A**). Overall, Abi-1-S296A mice showed a trend toward shorter survival than WT mice following WBI (**Fig. 5B**; 447 versus 565 days, respectively; p = 0.09). Approximately one year after irradiation, mice in the Abi-1-S296A group began to exhibit signs of morbidity. Three S296A mice which died prior to the study endpoint were excluded from tissue collection due to advanced postmortem decomposition. Notably, all three mice exhibited corneal erosion or ulceration. In one additional mouse, we were able to collect multiple organs and tissues, including the protruding and ulcerated eyes. Histopathological examination revealed the cause of the exophthalmos to be a papillary cystadenoma of the retrobulbar Harderian gland (**Table 2**).

**Figure 5.**
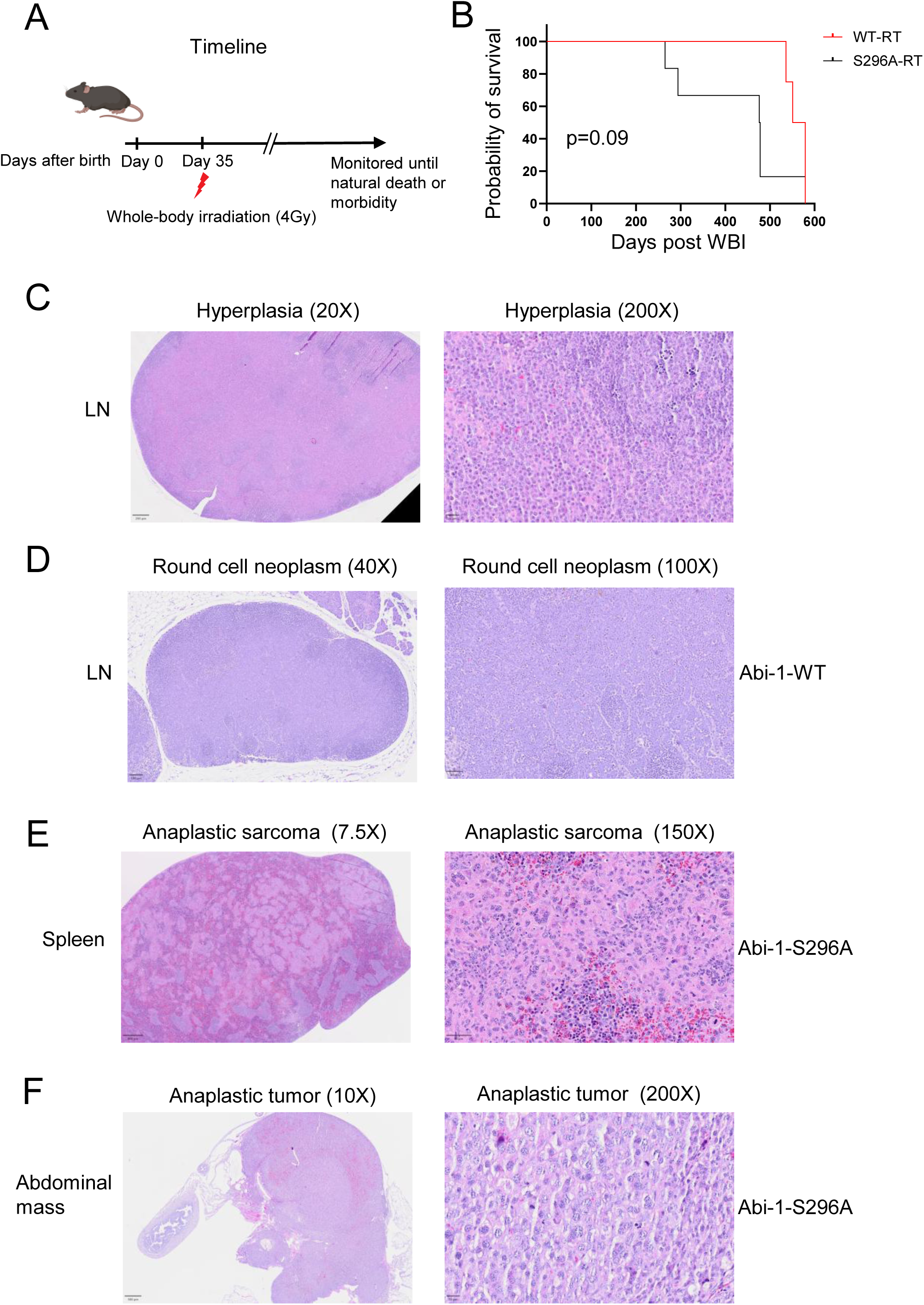
Abi-1-S296A mice show shorter survival and increased tumor development following low-dose whole-body irradiation. **(A)** Schematic timeline of the whole-body irradiation (WBI) experiment in WT and Abi-1-S296A knock-in mice. Mice received a single dose of 4 Gy at 5 weeks of age, followed by monitoring of their general health and survival. **(B)** Kaplan–Meier overall survival curves of irradiated WT and Abi-1-S296A mice. The p value was determined using the log-rank (Mantel–Cox) test. **(C)** Representative H&E-stained sections of reactive lymphoid hyperplasia from one of irradiated Abi-1-S323A mice were shown. **(D)** Representative H&E-stained sections of tumors collected from irradiated WT mice were shown. **(E-F)** Representative H&E-stained sections of tumors collected from irradiated Abi-1-S296A mice were shown. Tumor types and tissue origins are indicated in the corresponding images.

**Table 2.**
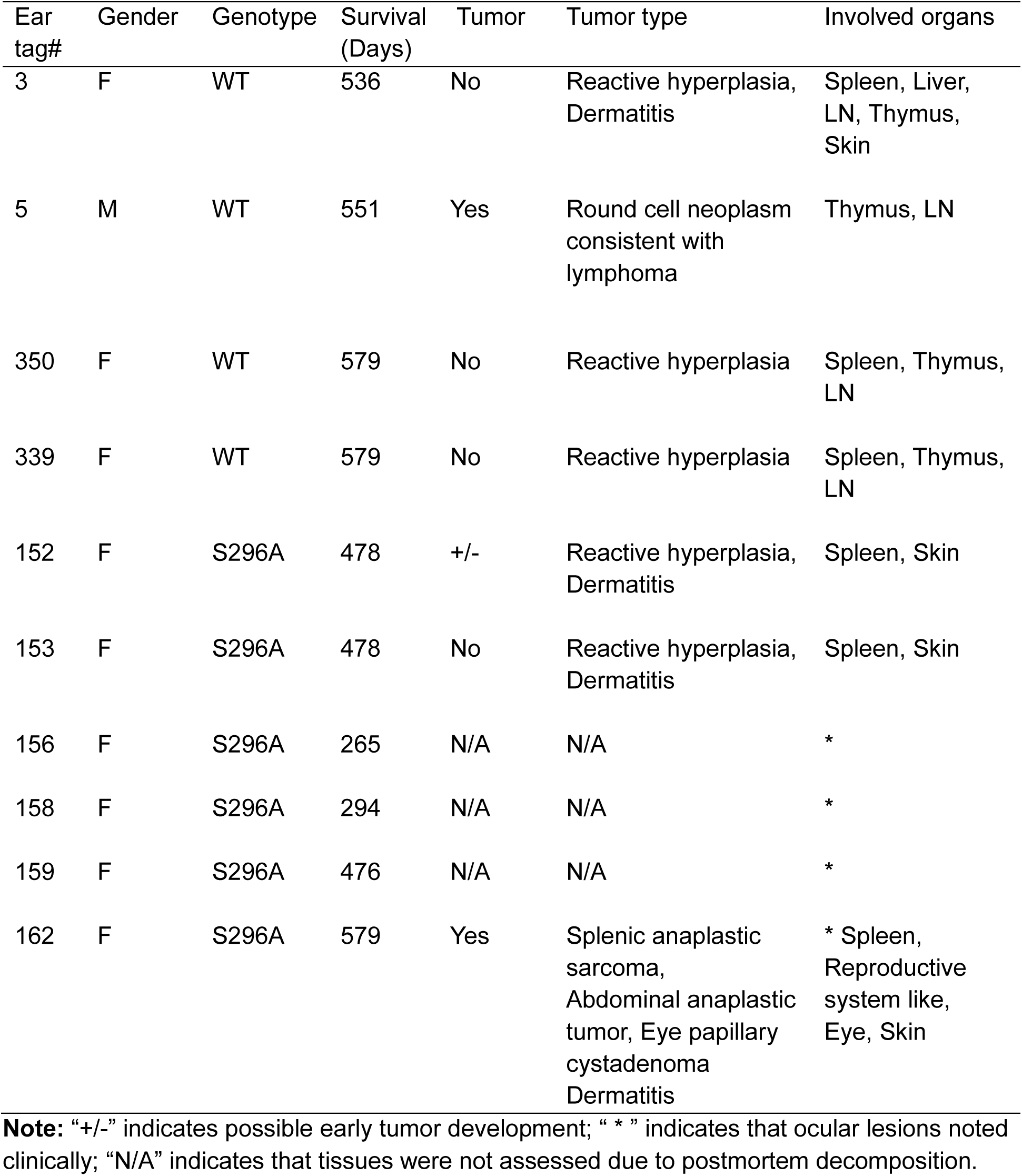
Survival Times and Tumor Types in Individual Mice Following Whole-Body Irradiation (4Gy)

| Ear tag# | Gender | Genotype | Survival (Days) | Tumor | Tumor type | Involved organs |
|---|---|---|---|---|---|---|
| 3 | F | WT | 536 | No | Reactive hyperplasia, Dermatitis | Spleen, Liver, LN, Thymus, Skin |
| 5 | M | WT | 551 | Yes | Round cell neoplasm consistent with lymphoma | Thymus, LN |
| 350 | F | WT | 579 | No | Reactive hyperplasia | Spleen, Thymus, LN |
| 339 | F | WT | 579 | No | Reactive hyperplasia | Spleen, Thymus, LN |
| 152 | F | S296A | 478 | +/- | Reactive hyperplasia, Dermatitis | Spleen, Skin |
| 153 | F | S296A | 478 | No | Reactive hyperplasia, Dermatitis | Spleen, Skin |
| 156 | F | S296A | 265 | N/A | N/A | * |
| 158 | F | S296A | 294 | N/A | N/A | * |
| 159 | F | S296A | 476 | N/A | N/A | * |
| 162 | F | S296A | 579 | Yes | Splenic anaplastic sarcoma,<br>Abdominal anaplastic tumor, Eye papillary cystadenoma<br>Dermatitis | * Spleen, Reproductive system like, Eye, Skin |

**Note:** “+/-” indicates possible early tumor development; “ \* ” indicates that ocular lesions noted clinically; “N/A” indicates that tissues were not assessed due to postmortem decomposition.

Lymphoid enlargement characterized primarily by reactive lymphoid and plasma cell hyperplasia was observed in both WT and S296A mice (**Fig. 5C**). Among the WT mice, one exhibited thymic and nodal round-cell neoplasia (**Fig. 5D**). In the S296A group, the mouse with the Harderian gland cystadenoma also exhibited two different malignant neoplasms. In the spleen the white pulp was largely effaced by a poorly differentiated anaplastic sarcoma (**Fig. 5E**). Then in the caudal abdomen, this same mouse also exhibited a separate poorly differentiated neoplasm morphologically distinct from the neoplasms identified in the spleen and harderian gland (**Fig. 5F**). Individual survival times and the tumor types identified in each mouse are summarized (**Table 2**).

### Abi-1-S296A promotes tumorigenesis in an AOM/DSS-induced colon cancer model

To further investigate whether Abi-1 dephosphorylation increases susceptibility to tumorigenesis following DNA damage, we employed an azoxymethane/dextran sodium sulfate (AOM/DSS)-induced colon tumorigenesis model. This approach was based on our previous finding that the Rac1-Abi-1-S323 signaling pathway is activated in intestinal tissues and promotes DNA repair, thereby alleviating intestinal injury [17]. The AOM/DSS model combines AOM-induced DNA damage with DSS-induced colonic epithelial injury and inflammation, providing a complementary approach to investigate tumor development in the context of DNA damage, tissue injury, and inflammation [35, 40, 41]. Following repeated cycles of AOM/DSS treatment, mouse colons were harvested approximately 80 days after AOM administration for tumor assessment (**Fig. 6A**). Compared with WT mice, S296A mice developed a greater number of tumors, which were also larger in size (**Fig. 6B–E**). These findings suggest that the Abi-1-S296A mutation increases susceptibility to AOM/DSS-induced colorectal tumorigenesis.

**Figure 6.**
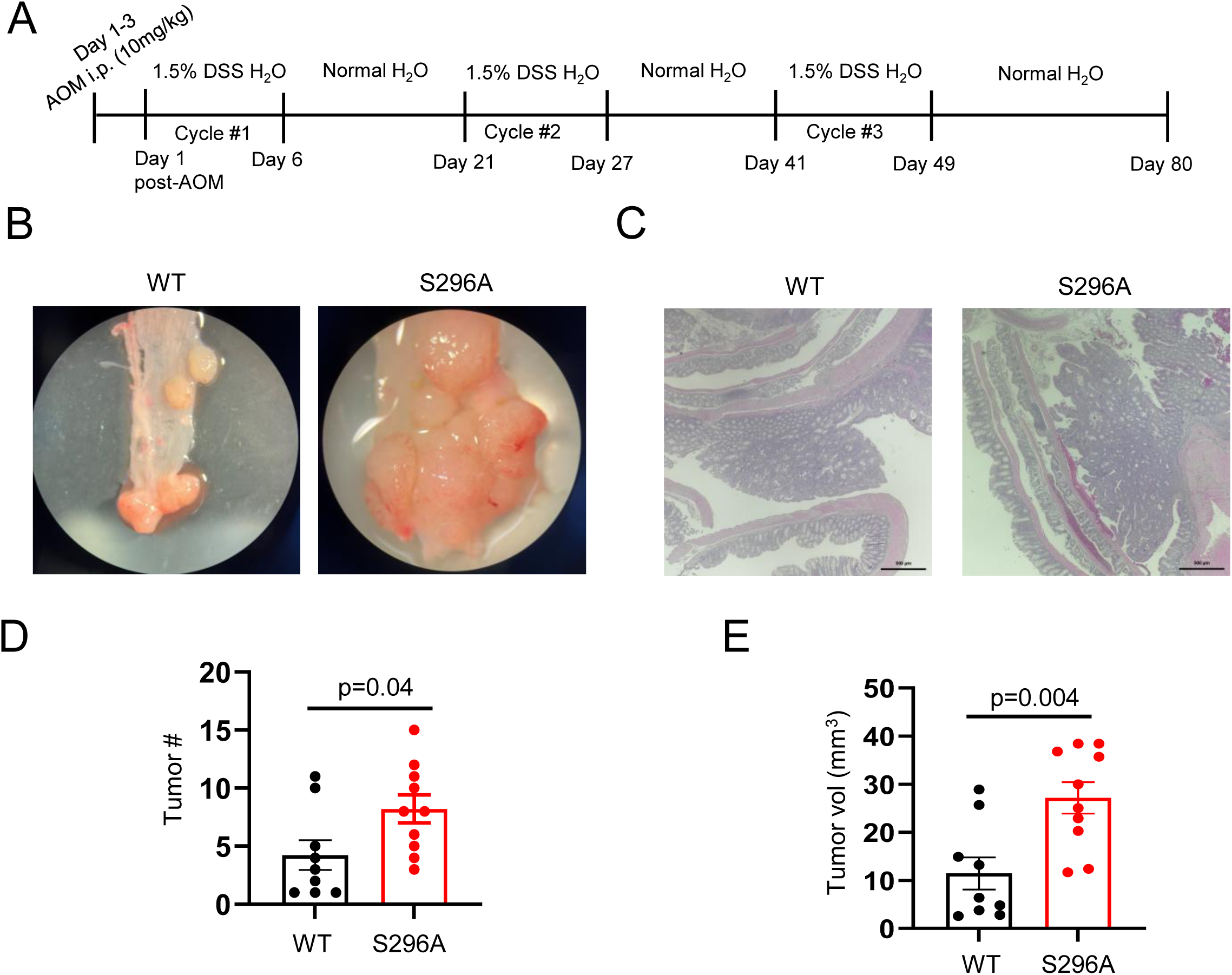
Abi-1 S296A promotes tumorigenesis in an AOM/DSS-induced colon cancer model. **(A**) Schematic timeline of azoxymethane/dextran sodium sulfate (AOM/DSS)-induced colon tumorigenesis in WT and Abi-1-S296A knock-in mice. Additional details of the treatment regimen are provided in Methods. **(B)** Representative gross images of colons showing tumors in WT and Abi-1-S296A mice. **(C)** Representative H&E-stained sections of colonic tumors from mice of the indicated genotypes. **(D)** Quantification of total tumor number per mouse in each group. Each dot represents one mouse (n = 10 mice per group). **(E)** Quantification of average tumor volume per mouse in each group. Each dot represents the average tumor volume for one mouse (n = 10 mice per group). Data are presented as mean ± SEM and were analyzed using a two-tailed Student’s t-test.

## Discussion

In this study, we demonstrate that Abi-1 S323 dephosphorylation promotes tumorigenesis across multiple nonmalignant cell models, an orthotopic tumor model, and multiple primary GEMMs, extending its previously established role in NHEJ-mediated DNA repair and treatment resistance [17]. Forced activation of this signaling pathway through expression of the dephospho-mimetic Abi-1-S323A mutant promotes transformation-associated phenotypes in nonmalignant cells and promotes orthotopic tumor growth under unperturbed conditions. Furthermore, using a newly generated CRISPR/Cas9-mediated Abi-1-S296A knock-in mouse model, in which murine S296A corresponds to human S323A that constitutively maintains the signaling active, we demonstrate that the non-phosphorylatable mutation at this site promotes spontaneous tumor development and increases susceptibility to tumorigenesis following DNA damage induced by whole-body irradiation or AOM/DSS treatment. These findings establish a previously unrecognized role for Rac1-Abi-1 signaling in tumorigenesis and highlight the utility of a site-specific Abi-1 knock-in mouse model for investigating the physiological and pathological consequences of this signaling.

Our findings challenge the conventional view that NHEJ protects genomic integrity by repairing DNA DSBs, highlighting the possibility that abnormal NHEJ activity may contribute to tumorigenesis. Although canonical NHEJ is generally considered an efficient and relatively precise pathway essential for DSB repair and genome maintenance, its ability to cause the incompatible DNA ends that may introduce mutations or deletions, causing genomic instability [5]. In this study, we found that activation of the Rac1-Abi-1 signaling pathway, which we previously established as a positive regulator of canonical NHEJ, promotes tumor development in two models of DNA damage-associated tumorigenesis (**Fig. 5 and 6**). Whole-body irradiation, which induces widespread DNA damage throughout the body, was associated with reactive hyperplasia and broad tumor development in multiple tissues and organs of S296A mice. In another model, AOM/DSS treatment induces localized DNA damage and inflammation in the colon, where the S296A mutation significantly enhanced colon tumorigenesis. These findings raise the possibility that Rac1-Abi-1 signaling promotes tumorigenesis in response to DNA damage, potentially through the abnormal NHEJ repair activity. However, the direct contribution of NHEJ to this process remains to be established. In addition, Rac1 signaling can activate downstream oncogenic pathways that modulate DNA repair and cellular responses to genotoxic stress [42, 43], which can potentially contribute to tumorigenesis. Loss or mutation of TP53 can increase Rac1 activity [44, 45], while PTEN loss has been associated with increased Abi-1 expression and enhanced Abi-1-dependent signaling[46]. Given the roles of TP53 and PTEN in maintaining genomic integrity in response to genotoxic stress, further studies are needed to determine the extent to which NHEJ dysregulation and crosstalk with other oncogenic and tumor-suppressive pathways contribute to the tumor phenotypes observed in Abi-1-S296A mice.

Our study establishes a new Abi-1-S296A knock-in mouse model that provides a unique tool for investigating the physiological and pathological functions of site-specific Abi-1 dephosphorylation. Conventional whole-body knockout of Abi-1 results in embryonic lethality [36], leading to the development of conditional and inducible Abi-1 knockout mouse models [37, 38]. However, these models eliminate the entire Abi-1 protein, potentially disrupting its other important cellular functions, including its role in maintaining cellular structure [47]. In contrast, our Abi-1-S296A model enables us to investigate the biological consequences of site-specific Abi-1 dephosphorylation without eliminating the protein or its other functions. Furthermore, this model provides a new approach to investigate the tumorigenic consequences of altered NHEJ signaling. Conventional mouse models used to investigate the role of NHEJ have targeted essential core NHEJ factors, such as DNA-PKcs [48], Ligase IV [49], Ku [50], or XRCC4 [51]. These genetic alterations disrupt fundamental DNA repair functions and can lead to severe developmental and immunological abnormalities. In contrast, our Abi-1-S296A model does not cause embryonic lethality and enables investigation of the biological consequences of constitutive activation of an upstream NHEJ-regulatory signaling pathway while preserving the core NHEJ machinery.

Our study also suggests that Abi-1 S296 phosphorylation plays important physiological role, as this mutation promotes spontaneous tumor development under unperturbed conditions. Notably, the spontaneous neoplasms observed in Abi-1-S296A mice predominantly involved lymphoid tissues and organs (**Fig. 4**). These findings raise the possibility that Rac1-Abi-1 signaling participates in regulating lymphocyte development or function. One important physiological process during lymphocyte development is V(D)J recombination, which generates antigen receptor diversity in developing B and T lymphocytes [52]. This process involves DNA DSBs introduced by the RAG1/RAG2 recombinase complex, which are subsequently repaired by classical NHEJ. Given that Abi-1 S323/296 dephosphorylation promotes NHEJ-mediated DSB repair, this pathway may also participate in the repair of RAG-induced DSBs during V(D)J recombination. Aberrant V(D)J recombination can result in abnormal antigen receptor rearrangements and chromosomal translocations, contributing to lymphoid malignancies [53, 54]. Thus, constitutive activation of Abi-1 signaling in Abi-1-S296A mice may alter the efficiency or fidelity of RAG-induced DNA end joining, potentially leading to abnormal antigen receptor rearrangements, chromosomal instability, and ultimately lymphoid tumorigenesis. Future studies examining V(D)J recombination, antigen receptor rearrangements, and the fidelity of RAG-induced DNA repair in Abi-1-S296A mice will help test this possibility.

Despite these findings, the mechanisms underlying Abi-1-S296A-associated tumorigenesis remain to be fully elucidated. Future studies will be important to determine whether dysregulated Rac1-Abi-1 signaling promotes tumor development by altering NHEJ repair fidelity following DNA damage or by affecting RAG-dependent V(D)J recombination and lymphocyte development under unperturbed conditions. In addition, investigating its interactions with other tumor-suppressive and oncogenic pathways may provide further insights into how Abi-1 S296 dephosphorylation contributes to tumorigenesis.

## Acknowledgments

Animal management, technical and experimental support services were provided by the University of Michigan Unit for Laboratory Animal Medicine (ULAM). E. McCulla was supported by the NCI Training Grant T32-CA009676, 1F31CA31406801. D.R. Wahl was supported by the NCI (R37CA258346), and the National Institute of Neurological Disorders and Stroke (R01NS129123). W. Zhou was supported by the UMMS Pandemic Research Recovery Program (U083054), Rogel Cancer Center Discovery Award and Michigan Radiosensitization SPORE Career Enhancement Program (5-P50-CA-269022-02). M. A. Morgan was supported by NCI (P50CA269022). L. D. Hess was supported by the NSF GRFP (DGE2241144). Research reported in this publication was partly supported by the National Cancer Institute of the National Institutes of Health under Award Number P30CA046592 through the use of the Rogel Cancer Center Experimental Irradiation Shared Resource and Cancer Data Science Shared Resource. The content is solely the responsibility of the authors and does not necessarily represent the official views of the National Institutes of Health. Research reported in this publication was supported by Rogel Comprehensive Cancer Center (P30CA046592) through the Experimental Irradiation Shared Resource. We acknowledge Zachary T Freeman, Thomas L. Saunders, Eden Dulka, Elizabeth Hughes, Hongali Zhang, Wanda Filipiak, and Galina Gavrilina and the Transgenic Animal Model Core (RRID:SCR_000776) of the University of Michigan’s Biomedical Research Core Facilities for design and production of the Abi knockin mice. Research reported in this publication was supported by the University of Michigan Transgenic Animal Model Core and the Biomedical Research Core Facilities. Research reported in this publication was supported by the National Cancer Institute under award number 5P30CA046592.

